# Virus-like particles (VLPs) are efficient tools for boosting mRNA-induced antibodies

**DOI:** 10.1101/2021.12.20.473421

**Authors:** Anne-Cathrine S. Vogt, Lukas Jörg, Byron Martina, Pascal S. Krenger, Xinyue Chang, Andris Zeltins, Monique Vogel, Mona O. Mohsen, Martin F. Bachmann

**Author notes:** equal contribution (Mona O. Mohsen, Martin F. Bachmann). Corresponding authors: Anne-Cathrine S. Vogt, Mona O. Mohsen.

## Abstract

mRNA based vaccines against COVID-19 have proven most successful at keeping the SARS-CoV-2 pandemic at bay in many countries. Recently, there is an increased interest in heterologous prime-boost vaccination strategies for COVID-19 to maintain antibody response for the control of continuously emerging SARS-CoV-2 variants of concern (VoCs) and to overcome other obstacles such as supply shortage, costs and reduced safety issues or inadequate induced immune-response. In this study, we investigate the antibody responses induced by heterologous prime-boost with vaccines based on mRNA and virus-like particles (VLPs). The VLP-based mCuMV_TT_-RBM vaccine candidate and the approved mRNA-1273 vaccine were used for this purpose. We find that homologous prime boost regimens with either mRNA or VLP induced high levels of high avidity antibodies. Optimal antibody responses were, however, induced by heterologous regimens both for priming with mRNA and boosting with VLP and vice versa, priming with VLP and boosting with mRNA. Thus, heterologous prime boost strategies may be able to optimize efficacy and economics of novel vaccine strategies.

## 1. Introduction

Multiple immunizations are usually required for most vaccines to be successful in protecting against a virus and its emerging variants. For instance, five-dose series of Tetanus, Pertussis and Diphtheria (DTaP) vaccine is required during childhood and an adolescent booster dose to elucidate the aimed protection (1). It is not entirely clear why some vaccines require more additional boosters than others; however, it is well accepted that multiple immunizations are essentials for better response, in particular for non-replicating vaccines (2). An annual dose of influenza vaccine is also recommended for persons who may be at increased risk for complications from influenza infections.

Additionally, heterologous prime-boost regimens in particular when using vectored vaccines have previously shown better immunogenicity; for example, priming with e.g. DNA and boosting with e.g. viral vectors (3, 4). The need to change the vector may be explained by the induction of neutralizing antibodies against the vector, which compromise further boosting with the same vector. Vaccine-specific antibodies have also been reported to suppress the cytotoxic (CTL) response when using the non-infectious virus-like particles (VLPs) derived from human papilloma virus (HPV) or Qβ-p33 for immunization (5, 6).

COVID-19 pandemic has overwhelmed the healthcare systems worldwide and several countries have prioritized the development of SARS-CoV-2 vaccine to contain the virus. Currently, 8 vaccines are approved in various countries and around 60 vaccines are in clinical development, of which 13 are in phase 3 (7). Several leading vaccines have been shown to confer protection and induction of neutralizing antibodies against the wild type SARS-CoV-2. However, with the continuous emergence of SARS-CoV-2 variants and the drop in antibody titers induced in vaccinated individuals, serious concerns are raised by public health organizations concerning the duration and efficacy of antibody responses by current vaccines (8).

Next-generation COVID-19 vaccines which have been approved for emergency use, namely mRNA vaccines such as BNT162b2 and mRNA-1273, have their own advantages and disadvantages. Obviously, the main advantage is their remarkable efficacy of 95% for Pfizer® mRNA vaccine and 94.1% for Moderna® (9, 10). On the other hand, their adverse reactions profile has raised concerns with respect to their safety profile. The incidence of local and systemic adverse reactions (AR) has been shown to be relatively high for mRNA and adenovirus-based vaccines with Local AR reaching 40%-88.9% and systemic AR between 44% and 86%, respectively (11).

Virus-like particles (VLPs) are considered traditional vaccine platforms. They are virus-derived structures and have the ability self-assemble to mimic the parental virus in size and shape. One important aspect is that VLPs lack any genetic materials; accordingly, they are not capable of infecting the host cell or replication (12). Such traditional VLP-based vaccines have been approved for human use against different viruses decades ago. For example, vaccines against Hepatitis B and E viruses (HBV and HEV), Human Papilloma Virus (HPV) and the newly developed malaria vaccine (13). VLP-based vaccines have also been tested for COVID-19 in preclinical and clinical trials. The traditional vaccine platforms have shown lower incidence of adverse side effects (14); however they typically also showed inferior immunogenicity in comparison to the next-generation mRNA vaccines (15, 16).

In line with the above, heterologous prime-boost vaccination strategies using different vectors should be tested to elicit broader and more efficient protective immune-responses with better and improved safety profiles. This strategy may meet the emergency needs in the current pandemic. In the current study, we evaluated different prime and boost regimens combining mRNA-1273 with a VLP-based vaccine based, namely mCuMV_TT_-RBM. We have previously designed and developed the scalable and immunogenic VLP-based COVID-19 vaccine mCuMV_TT_-RBM, which has been shown to induce RBD-specific IgG and IgA antibodies with strong neutralizing capability, albeit inferior to mRNA induced response (15). We show that a booster dose with mRNA-1273 or mCuMV_TT_-RBM induced high levels of high avidity antibodies. Interestingly, highest antibody responses were measured, when vaccination was performed by heterologous administration of mRNA for prime and VLP as boost and vice versa, priming with VLP and boosting with mRNA. Along these findings, heterologous prime boost strategies may be able to improve efficacy and economics of current vaccine protocols.

## 2. Results

### 2.1. Heterologous prime-boost vaccine administration induces comparable high levels of RBD-specific antibodies following the booster dose

We have designed different heterologous vaccination regimens using mRNA-1273 vaccine (Moderna®) and our newly developed mosaic COVID-19 VLPs-based vaccine mCuMV_TT_-RBM (15) as illustrated in Figure 1A. For SARS-CoV-2 specific vaccine, the mRNA provides the genetic expression of the spike protein of COVID-19 (17). The mRNA of the spike protein is then translated by the host, thus allowing the host to mount an antibody response against the expressed protein (18, 19). mRNA-1273 has shown >90% effectiveness in preventing the SARS-CoV-2 infection, at least for the original Wuhan strain (10). mCuMV_TT_-RBM is a plant-derived VLPs which incorporates the receptor-binding motif (RBM) of SARS-CoV-2 using genetic fusion techniques. mCuMV_TT_-RBM also incorporates in its interior surface a tetanus toxin (TT) epitope which is believed to enhance the immune response in elderly people. Specifically, the TT-epitope is expected to augment the interaction between TT-specific T helper (T_H_) cells and RMB-specific B cells. This is supported by the fact that pre-existing immunity to the chosen TT epitope is very broad in humans (and animals) as the peptide binds essentially to all HLA-DR molecules and most people have been immunized many times against TT. Additionally, mCuMV_TT_-RBM is packaged with ssRNA, a TLR7/8 ligand and serves as a natural adjuvant (20, 21).

**Figure 1.**
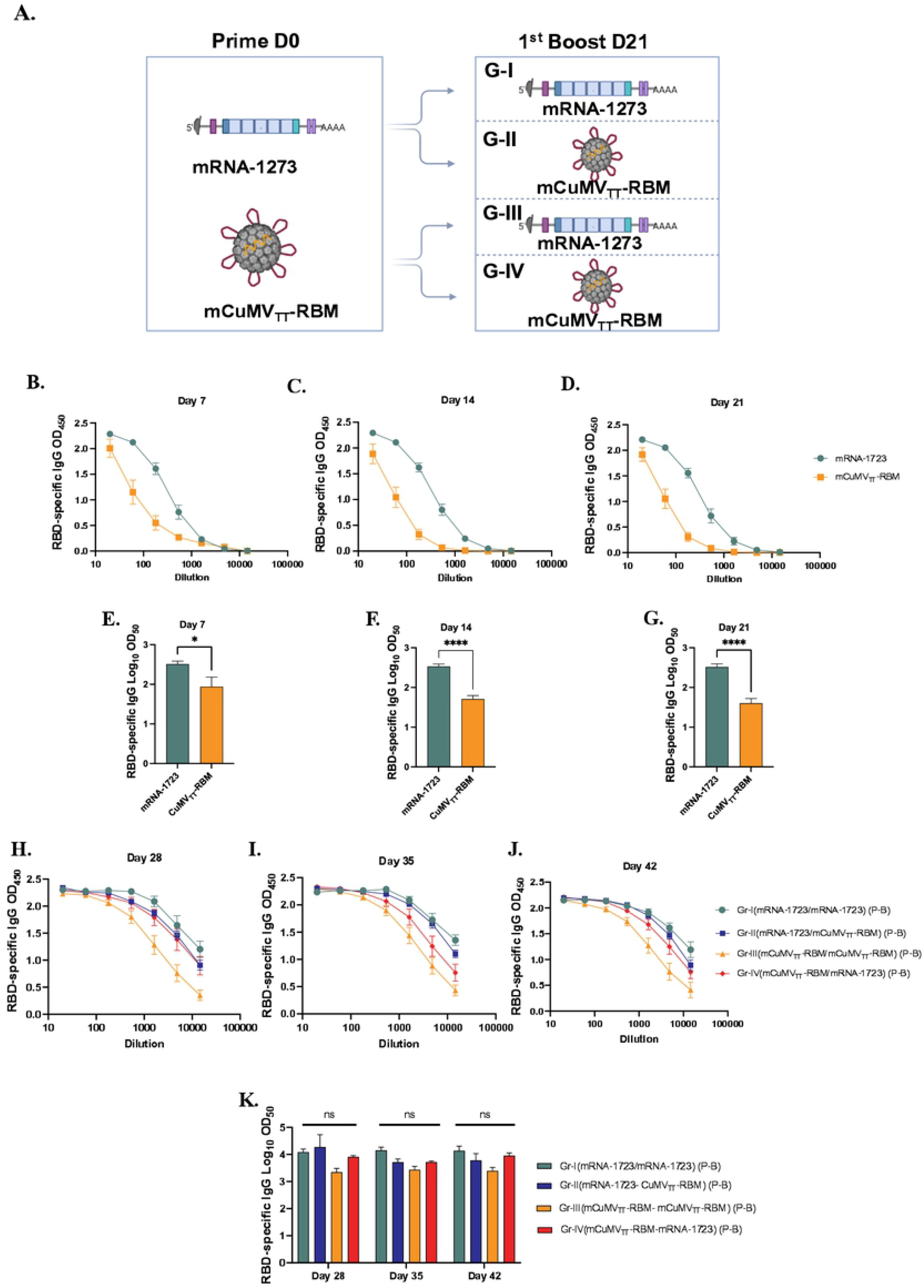
Heterologous prime-boost vaccine administration induces comparable high levels of RBD-specific antibodies following the booster dose. Heterologous prime boost vaccination induces high levels of RBD-specific antibodies. A) Vaccination regiment (Prime/Boost) D0/D21 and groups. B-D) OD_450_ of RBD-specific IgG for the groups vaccinated with mRNA-1723 or mCuMV_TT_-RBM on days 7, 14 and 21. E-G) Log_10_OD_50_ of RBD-specific IgG titers for the groups vaccinated with mRNA-1723 or mCuMV_TT_-RBM on days 7, 14 and 21. H-J) OD_450_ of RBD-specific IgG for the groups boosted with mRNA-1723 or mCuMV_TT_-RBM on days 28, 35 and 42. K) Log_10_OD_50_ of RBD-specific IgG titers for the groups boosted with mRNA-1723 or mCuMV_TT_-RBM on days 28, 35 and 42 using D0/D21 regimen. Statistical analysis (mean ± SEM) using one-way ANOVA in K or Student’s t test in E-G, n=10 or 5. One representative of 3 similar experiments is shown. The value of p<0.05 was considered statistically significant (*p<0.01, **p<0.001, ***p<0.0001).

10μg/dose of mRNA-1723 vaccine and 100μg/dose of mCuMV_TT_-RBM were used in this experiment for priming and boosting. Balb/c mice were divided into two groups for subcutaneous (s.c.) priming with mRNA-1723 or mCuMV_TT_-RBM vaccines (Fig. 1A). Blood samples were collected on a weekly basis to measure the induced RBD-specific antibodies by ELISA. Our results showed successful induction of RBD-specific antibodies 7 days after subcutaneous injection of mRNA-1723 or mCuMV_TT_-RBM (Fig. 1B). mRNA-1723 was superior to mCuMV_TT_-RBM in inducing RBD-specific antibodies measured on days 7, 14 and 21 (Fig.1C-G) but the VLPs also was effective at inducing rapid IgG responses.

The primed mice were divided next into four groups (Fig. 1A) and received a booster dose on day 21 using mRNA-1723 or mCuMV_TT_-RBM. Therefore, mice are grouped and immunized as follows: Group I (mRNA-1723 → mRNA-1723), Group II (mRNA-1723 → mCuMV_TT_-RBM), Group III (mCuMV_TT_-RBM → mCuMV_TT_-RBM) and Group IV (mCuMV_TT_-RBM → mRNA-1723). Blood samples were collected on days 28 and 35 to measure the induced RBD-specific antibodies by ELISA. Interestingly, heterologous boost with mRNA-1723 or mCuMV_TT_-RBM showed a similar induction of RBD-specific antibodies with no statistical difference for sera collected on days 28, 35 and 42 (Fig. 1H-K). This demonstrates that mCuMV_TT_-RBM can significantly boost previously primed B-cells to a similar titer when using a homologous mRNA vaccine.

### 2.2. A second booster dose further enhances the induced immune response

Taking into account the slightly inferior immunogenicity of mCuMV_TT_-RBM vaccine to mRNA-1723, and our previous data showing that a 2^nd^ booster dose of mCuMV_TT_-RBM would enhance the quality of the induced antibodies (15), a 2^nd^ booster dose was performed on day 56 in Groups II, III and IV as illustrated in Figure 2A. As Group I represents the standard immunization strategy followed in almost all countries, no booster dose was applied. ELISA data for OD450 and OD50 for days 63 and 70 did not show an increase in RBD-specific antibodies in comparison to the induced antibodies following the 1^st^ boost (Fig. 2B-D).

**Figure 2.**
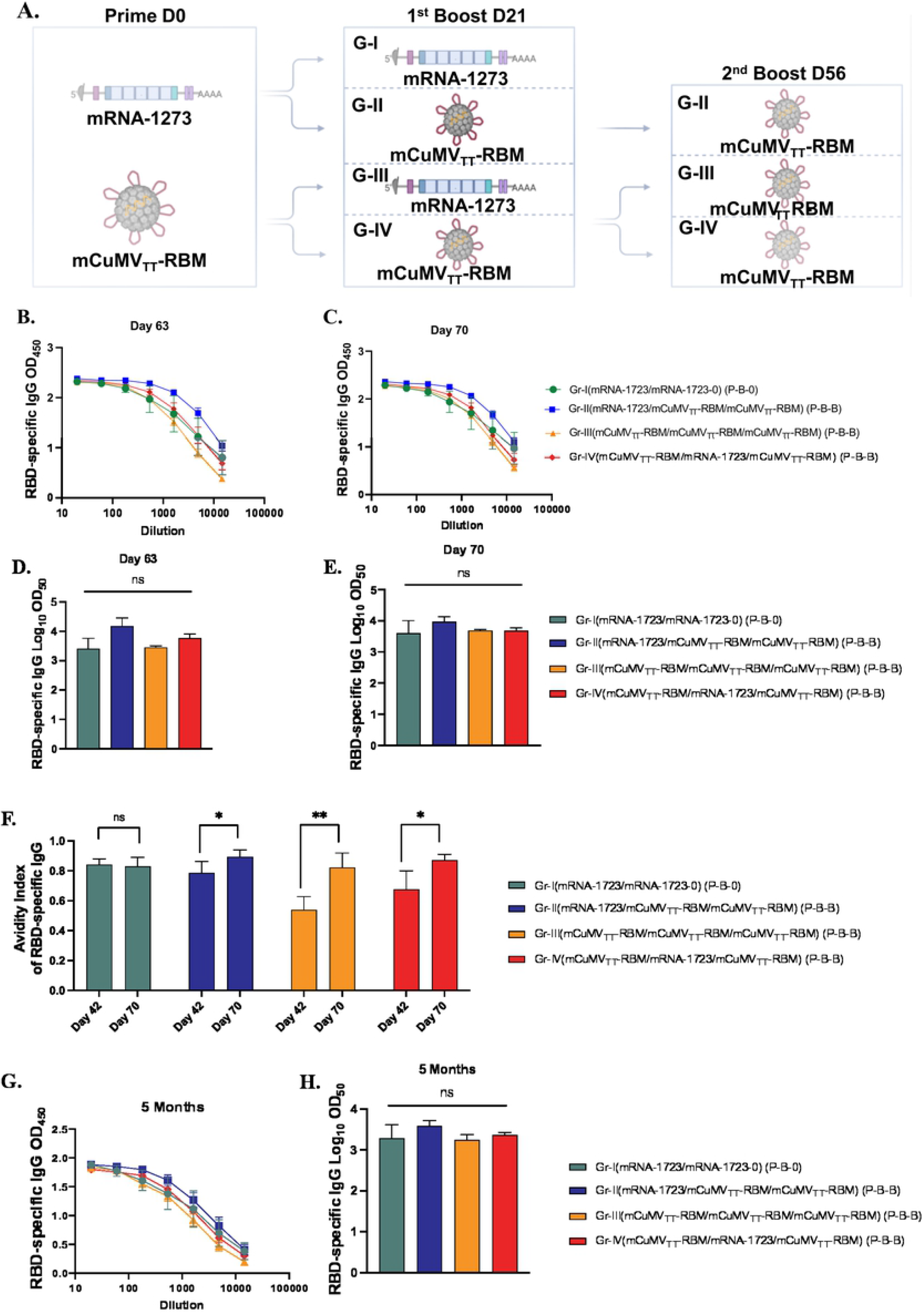
A second booster dose further enhances the induced immune response. Enhanced immune response with second booster dose. A) Vaccination regiment (Prime-Boost-Boost) D0/D21/D56 and groups. B and C) OD_450_ of RBD-specific IgG for the groups vaccinated with mRNA-1723 or mCuMV_TT_-RBM on days 63 and 70 using D0/D21/D56 regimen. D and E) Log_10_OD_50_ of RBD-specific IgG titers for the groups vaccinated with mRNA-1723 or mCuMV_TT_-RBM on 63 and 70 using D0/D21/D56 regimen. F) Avidity Index of RBD-specific IgG in mice vaccinated with mRNA-1723 or mCuMV_TT_-RBM using D0/D21 or D0/D21/D56 regimens, sera were treated with PBST or 7 M Urea. G) OD450 of RBD-specific longeivity IgG for the groups vaccinated with mRNA-1723 or mCuMV_TT_-RBM 5 months after priming using D0/D21/D56 regimen. H) Log_10_OD_50_ of RBD-specific IgG titers for the groups vaccinated with mRNA-1723 or mCuMV_TT_-RBM 5 month after priming, using D0/D21/D56 regimen. Statistical analysis (mean ± SEM) using one-way ANOVA in D,E and H or Student’s t test in F, n=10 or 5. One representative of 3 similar experiments is shown. The value of *p<0.05 was considered statistically significant (**p<0.01).

The avidity of the induced antibodies was assessed using a modified immunoassay using 7M urea which facilitates the detachment of the low avidity antibodies. More specifically, we compared the avidity of RBD-specific antibodies on days 42 and 70, i.e. after 2 or 3 injections. For group I, which was primed and boosted with mRNA-1723, the avidity of RBD-specific antibodies didn’t differ on days 42 and 70 (p. 0.7460). In Group II (which received a prime with mRNA-1723 and 2 booster doses of mCuMV_TT_-RBM) ~80% of RBD-specific antibodies were of high avidity after the 1^st^ boost in comparison to ~90% following the 2nd booster dose (*p*. 0.0436). This finding was similar in group IV (which received a prime with mCuMV_TT_-RBM, followed by a 1^st^ boost with mRNA-1723 and 2^nd^ boost with mCuMV_TT_-RBM), again with statistically significant difference in avidity between day 42 and 70 (*p*. 0.0243) (Fig. 2F). The induction of higher avidity antibodies following 3 vaccinations with mCuMV_TT_-RBM is consistent with our previous findings (15).

It was also of interest to study the longevity of the induced antibodies after vaccination with the different heterologous strategies. Accordingly, we have tested RBD-specific antibodies up to 5 months after the priming dose. The results revealed a slight drop in the antibody titer in group I (received 2 doses of mRNA-1723), however was not significantly different when compared to the other groups. Groups II, III and IV showed a stable antibody titer comparable to the titers seen on days 42 and 70 (Fig. 2G and H).

### 2.3. The induced antibodies recognize SARS-CoV-2 variants of concern (VoCs) efficiently

To test the capacity of the induced antibodies following the heterologous prime-boost regimen to neutralize SARS-CoV-2 wild-type and its mutated delta variant, we used a reduction of cytopathic effect (CPE) assay. 100 TCID_50_ of SARS-CoV-2/ABS/NL20 or delta strain have been used and titers have been expressed as the highest dilution that inhibits 50% CPE formation. No significant difference was detected between the four groups when measuring the neutralization titer against SARS-CoV-2 wild-type (Fig.3A). When comparing the induced neutralization titer against the delta VoC, our results show a significant difference between Group I (2xRNA) and III (3xVLP) (*p*. 0.039) confirming the superiority of mRNA only over VLP-only based vaccines. However, no statistical differences have been detected between Group I and II (*p*. 0.2558) or between Group I and IV (*p* >0.999) (Fig. 3B). This indicates that mRNA may be used to boost VLP-induced antibody responses; or vice versa, and probably more importantly, VLPs may be used to boost RNA-induced responses. Hence, classical VLP-based vaccines may be used to boost mRNA induced antibody responses.

**Figure 3.**
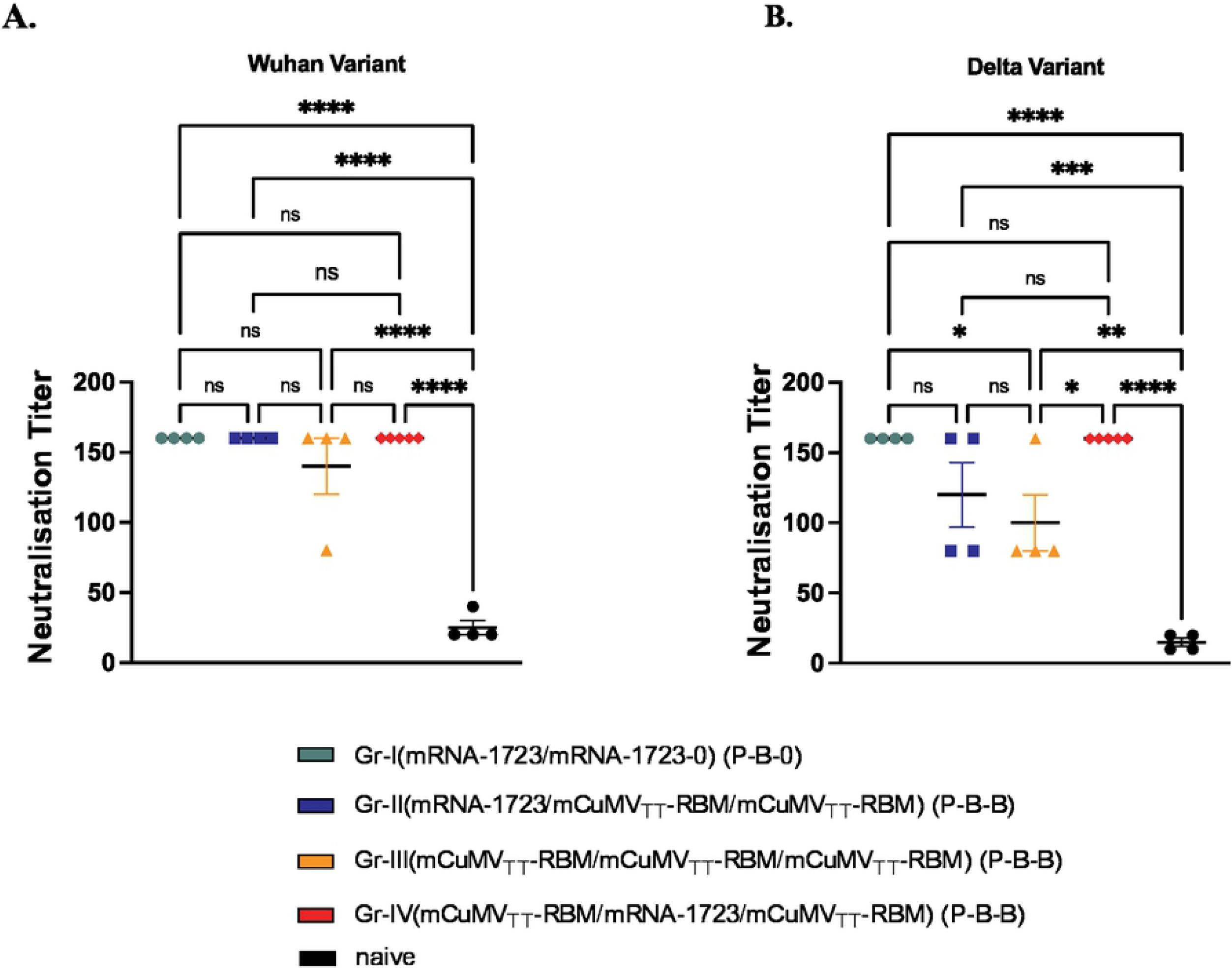
The induced antibodies recognize SARS-CoV-2 variants of concern (VoCs) efficiently. Recognition of SARS-CoV-2 variants of concern. A and B) Neutralization titer (CPE) for sera from mice vaccinated with mRNA-1723 and mice vaccinated with mCuMV_TT_-RBM, sera from day 63 (after 2nd boost). Statistical analysis (mean ±SEM) using ANOVA, n=5. One representative of 2 similar experiments is shown. The value of *p<0.05 was considered statistically significant (**p<0.01, ***p<0.001, ****p<0.0001).

## 3. Discussion

The highly contagious SARS-CoV-2 virus has put a heavy toll on the world’s public health systems and caused a disruption of the world economy (22). The use of masks, physical distancing, contact tracing as well as isolation of sick or infected people are important but insufficient measures to reduce the spread of the corona virus disease (23). Accordingly, vaccine campaigns are needed to curb the spread of SARS-CoV-2 and thereby reducing the mortality and morbidity associated with the disease.

The commercially available mRNA vaccines require stringent storage conditions during transport and at vaccination sites as they have a short half-life (24). According to manufactures information, mRNA-1273 is stable up o 6 month when kept at −20°C, whereas to store BNT162b2 for the same time-period, temperatures between −80°C to −60°C are needed.^9,10^ With regard to stability at room temperature, mRNA-1273 is stable for up to 12h; in contrast BNT162b2 has to be administered within 6h after thawing (25). These strict storage conditions represent a major obstacle for efficient use in less affluent parts of the world. Beside the challenging storage conditions, a further obstacle represent the production-costs of RNA-based vaccines, limiting their global access. BNT162b2 has a production cost of $19.50 and mRNA-1273 with even higher costs at $32-37 (11, 26). This can cause a major problem in low-income countries, to afford these types of vaccines. Therefore, cost effective and efficient vaccine candidates are of high importance taking into account the continuous persistence of the virus and the obstacles in eradicating it worldwide. We have previously shown high stability of our newly developed VLP-based vaccine-candidate mCuMV_TT_-RBM which was stable for at least 14 months at 4°C without signs of degradation (15). This has also been confirmed with other preclinical studies showing the stability of mCuMV_TT_-based vaccines for up to 12 months when kept at 4°C. Thus indicating a significant advantage for such VLP-based vaccines (27). Additionally, the production of mCuMV_TT_-RBM is highly scalable allowing the production of millions of the doses in a single 1000L fermenter run (15). The rapid pace of mRNA vaccine development accompanied fears of potential long-term adverse effects have raised some concerns in global community. VLP-based vaccines are therefore of high interest as they display a safe platform, due to the lack of replicating genetic material which is also been confirmed in the approved and marketed vaccines (HBV-HEV-HPV and recently Mosquirix™) (13). In addition, the here employed mCuMV_TT_ technology does not need use of an adjuvant.

Our data show that both vaccine-types, mRNA-1723 and mCuMV_TT_-RBM are capable of inducing high RBD-specific antibody titers 7 days following the priming dose. Interestingly, mRNA-1723 shows a superior induction of antibody response in comparison to the VLP-based vaccine. However, after the administration of the 1^st^ homologous boosting, this difference was not statistically significant anymore (Group I and III). The same was true heterologous boosts (Group II and IV), indicating the ability of VLPs to efficiently boost the B cells primed by mRNA. We have previously hypothesized that natural infection with SARS-CoV-2 virus induces short-lived neutralizing antibodies due to the unusually large distance between RBD-epitopes embedded in the membrane surface (28). This obstacle may be overcome by grafting RBM or RBD onto highly repetitive nanoparticles such as the here used mCuMV_TT_ VLPs, resulting in successful display of the virus-epitopes at optimal distances of 5-10nm. This pathogen-associated structural pattern (PASP) may also play a role in the induction of higher avidity antibodies as shown after the 2^nd^ booster dose.

With the continuous emergence of SARS-CoV-2 variants (VoCs), it may be essential to administer repetitive booster doses, as higher antibody levels may need to be maintained for sustained protection. Indeed, our data demonstrate that a 2^nd^ boost can maintain antibody titers. Furthermore, the 2^nd^ boost in the different heterologous regimens in groups II, III and IV have significantly enhanced the avidity of the antibodies, likely a key attribute for neutralization of emerging new variants.

The neutralizing capacity of the induced antibodies following the homologous or heterologous vaccination strategies showed that both approaches are efficient against the wild-type SARS-CoV-2. No statistical difference in the ability to neutralize the delta variant has been detected between groups I, II and IV. Furthermore, heterologous boosting of mice primed with mRNA-1723 or mCuMV_TT_-RBM revealed similar outcome. These data confirm the ability of mCuMV_TT_-RBM to efficiently boost antibodies primed with mRNA-1723 vaccine in a similar way to the standard followed vaccination strategy (mRNA-1723 prime/ boost).

Collectively, our data support the premise of using a heterologous prime/boost vaccination regimen against COVID-19. The CuMV_TT_-RBM VLPs may therefore constitute an efficient platform for boosting previously primed B cell responses; here we demonstrate this for mRNA primed responses, but it is likely that this may extend to B cell responses primed by other modalities, including vector- or virus induced immune reactions.

## 4. Materials and Methods

### 4.1. Mice

*In vivo* experiments were performed using 8-12 weeks-old female, BALB/cOlaHsd mice purchased from Envigo (Amsterdam, Netherlands). All mice were maintained in microisolater cages with free access to water and food. Mice were kept on standard chow diet ((diet Catalog # 3430); Granovit AG-Kliba Nafag, Switzerland). All animals could acclimatize to the facility for one week before experiments were performed. All animal procedures were conducted in accordance with the Swiss Animal Act (455.109.1 – September 2008, 5^th^) of University of Bern. All animals were treated for experimentation according to protocols approved by the Swiss Federal Veterinary Office.

### 4.2. Vaccines

mRNA-1273 (Moderna®) was kindly provided the Inselspital, Bern University Hospital. Mosaic (mCuMV_TT_-RBM) vaccine was prepared as previously described by Mohsen *et al.* (15).

### 4.3. Vaccination regimen/dose/sera collection

Wild type BALB/cOlaHsd mice were vaccinated subcutaneously (s.c.) using different regimens as summarized in Table 1. 10μg (Prime on day 0) and 10μg for a booster dose on day 21 were used. mCuMV_TT_-RBM vaccine and mRNA-1273 were diluted in 1xPBS in a final volume of 100μl for final injection. Serum was collected on a weekly basis. The three doses regimen (prime-1^st^ boost-2^nd^ boost) was either 10μg prime and 100μg 1^st^ and 2^nd^ boost or 100μg prime and 10μg 1^st^ boost and 100mg 2^nd^ boost given at days 0, 21 and 56.

**Table 1:**
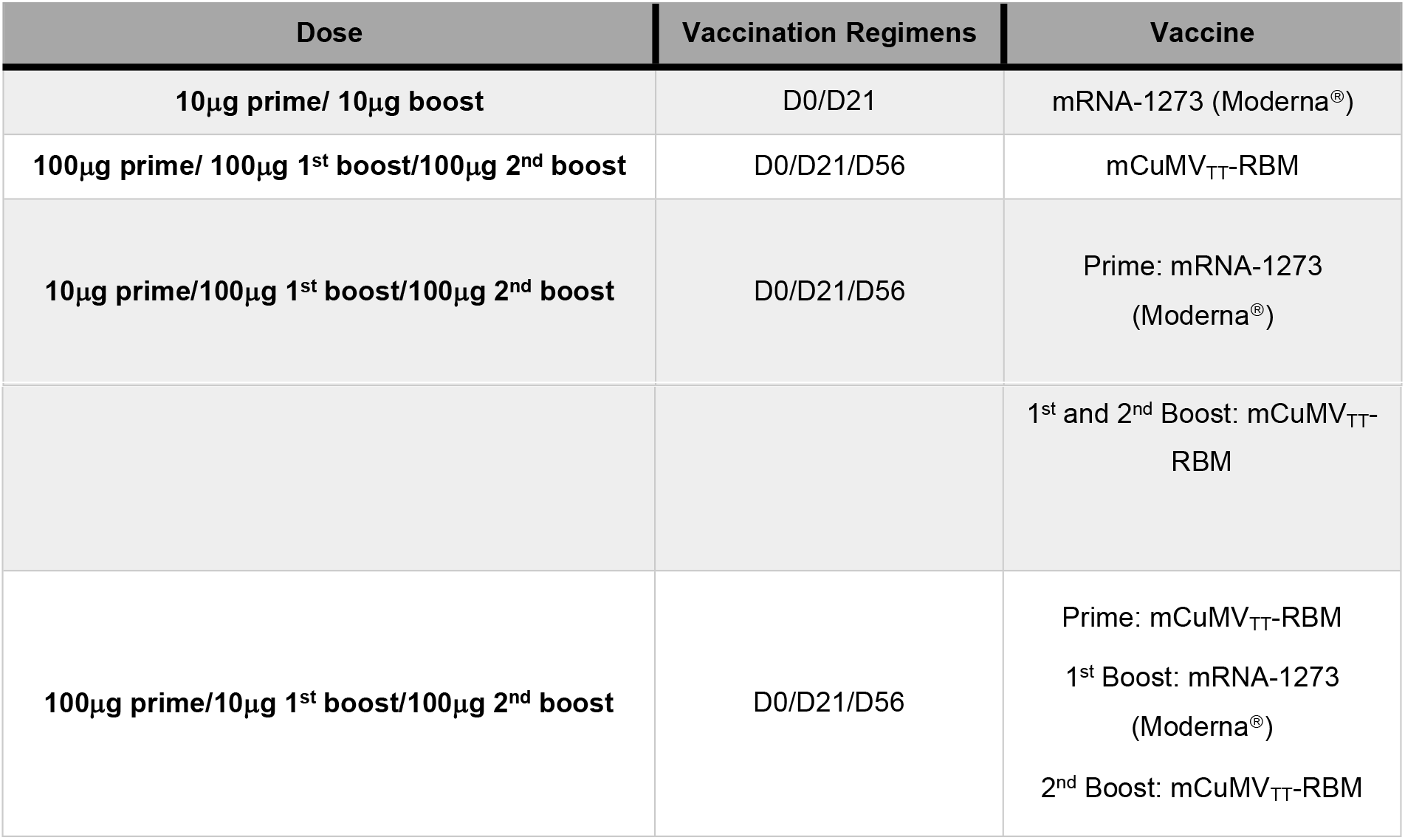
Used Doses and Vaccine Regimens

### 4.4. Expression and purification of RBD

SARS-CoV-2 RBD_wildtype_, was cloned as a synthetic gene into pTWIST-CMV-BetaGlobin-WPRE-Neo vector (Twist Biosciences, CA, USA) and expressed in HEK293F cells through the Expi293 system (ThermoFisher Scientific, MA, USA). Purification was performed by IMAC using a HiTrap TALON crude column (Cytiva, Uppsala, Sweden).

### 4.5. Enzyme-linked immunosorbant assay (ELISA)

To determine the total IgG antibodies against the vaccine mCuMV_TT_-RBM as well as mRNA-1273 in sera of vaccinated mice, ELISA plates (96 well half-area ELISA plates; Costar, Corning, Catalog # 3690) were coated with SARS-CoV-2 RBD (wildtype) at a concentration of 1μg/ml overnight at 4°C. ELISA plates were washed with PBS-0.01% Tween and blocked using 100μl PBS-Casein 0.15% for 2h in RT. Sera from vaccinated mice was serially diluted 1:3 starting with a dilution of 1:20 and incubated for 2h at RT. After washing with PBS-0.01%Tween, goat anti-mouse IgG conjugated to Horseradish Peroxidase (HRP) (Jackson ImmunoResearch, West Grove, Pennsylvania) was added at 1/1000 and incubated for 1h at RT ELISA was developed with tetramethylbenzidine (TMB), stopped by adding equal 1 M H2SO4 solution. OD450 was measured using the SpectraMax M5 ELISA reader (Molecular Devices, Catalog # M5) and read at OD_450_ nm or expressed as Log OD_50_. Detecting RBD-specific IgGs against mutated RBDs was carried out in a similar way.

### 4.6. Avidity (ELISA)

To test IgG antibody avidity against RBD protein, threefold serial dilutions of 1/20 diluted mice sera, were added to ELISA plates (96 well half-area ELISA plates; Costar, Corning, Catalog # 3690) coated over night at 4°C with 1μg/ml RBD. After incubation at RT for 1h, the plates were washed once in PBS-0.01% Tween, and then washed 3x with 7M urea in PBS-0.05%Tween or with PBS-0.05% Tween for 5min every time. After washing with PBS-0.05%Tween, goat anti-mouse IgG conjugated to Horseradish Peroxidase (HRP) (Jackson ImmunoResearch, West Grove, Pennsylvania) was added 1/1000 and incubated for 1h at RT. Plates were developed and read at OD450 nm.

### 4.7. Cytopathic effect-based neutralization assay (CPE)

To determine the neutralizing ability and capacity of vaccine induced antibodies a CPE assay was performed using wild-type SARS-CoV-2 (SARS-CoV-2/ABS/NL20) and delta Strain. Serum samples were heat-inactivated for 30min at 56°C. Two-fold serial dilutions were prepared starting at 1:20 up to 1:160. 100 TCID_50_ of the virus was added to each well and incubated for 37°C for 1h. The mixture has been added on a monolayer of Vero cells and incubated again for 37°C for 4 days. Four days later the cells were inspected for cytopathic effect. The titer was expressed as the highest dilution that fully inhibits formation of CPE.

### 4.8. Statistical analysis

All data are presented as mean ± SEM. Data were analyzed using Ordinary One-way ANOVA for multiple comparisons and *Students’ t-test* when comparing two groups. At least two independent experiments were performed. Statistical significance was set at p≤0.05. *P< 0.05, **P < 0.01, ***P <0.001, ****P <0.0001. Analyses were performed using GraphPad PRISM 9.0 (Graph-Pad Software, Inc., La Jolla, CA, USA).

## ACKNOWLEDGMENT

This work was supported by Saiba AG and Inselspital Bern.

## AUTHORS’ CONTRIBUTIONS

ASV, LJ, PSK, XC, AZ, MOM, MFB: Design of experiments, acquisition of data, interpretation, and analysis of data. ASV, MOM, MV, MFB: Writing, revision, and editing of manuscript. LJ, MV, AZ: Technical, material, and tool support. MOM and MFB: Study supervision. All authors read and approved the final manuscript.

## DATA AVAILABILITY STATEMENT

The datasets generated during and/or analyzed during the current study are available from the corresponding author on reasonable request.

